# The Structural Layers of the Porcine Iris Exhibit Inherently Different Biomechanical Properties

**DOI:** 10.1101/2022.12.11.519999

**Authors:** Royston K. Y. Tan, Satish K. Panda, Fabian A. Braeu, Arumugam R. Muralidharan, Monisha E. Nongpiur, Anita S. Y. Chan, Tin Aung, Raymond P. Najjar, Michaël J.A. Girard

## Abstract

**Purpose:** To isolate the structural components of the *ex vivo* porcine iris tissue and to determine their biomechanical properties.

**Methods:** The porcine stroma and dilator tissues were separated, and their dimensions were assessed using optical coherence tomography (OCT). The stroma underwent flow test (*n* = 32) to evaluate for permeability using Darcy’s Law (Δ*P* = 2000 Pa, *A* = 0.0391 mm^2^), and both tissues underwent stress relaxation experiments (ε = 0.5 with initial ramp of δε = 0.1) to evaluate for their viscoelastic behaviours (*n* = 28). Viscoelasticity was characterised by the parameters *β* (half width of the Gaussian distribution), τ_*m*_(mean relaxation time constant), *E*_0_ (instantaneous modulus) and *E*_∞_ (equilibrium modulus).

**Results:** For the stroma, the hydraulic permeability was 9.49 ± 3.05 × 10^-6^ mm^2^/Pa·s, and the viscoelastic parameters were *β* = 2.50 ± 1.40, and τ_*m*_ = 7.43 ± 4.96 s, with the two moduli calculated to be *E*_0_= 14.14 ± 6.44 kPa and *E*_∞_ = 6.08 ± 2.74 kPa. For the dilator tissue, the viscoelastic parameters were *β* = 2.06 ± 1.33 and τ_*m*_ = 1.28 ± 1.27 s, with the two moduli calculated to be *E*_0_ = 9.16 ± 3.03 kPa and *E*_∞_ = 5.54 ± 1.98 kPa.

**Conclusion:** We have established a new protocol to evaluate the biomechanical properties of the structural layers of the iris. Overall, the stroma was permeable and exhibited smaller moduli than those of the dilator muscle. An improved characterisation of iris biomechanics may form the basis to further our understanding of angle closure glaucoma.

## Introduction

Research in angle closure glaucoma explores a myriad of possible risk factors and characteristics that are associated with the disease, ranging from genetic studies^1–5^ to anatomical measurements^6–10^ and computational studies^11–15^. Being able to predict whether a patient will be afflicted with angle closure is paramount, so that preventive measures can be undertaken^16, 17^. Similarly, clinical examinations can identify patients with a higher risk of developing angle closure so that laser iridotomy can be done to widen the drainage angle, lowering the chance of glaucoma development in many instances.

Whilst clinical examinations are crucial to aid early angle closure detection^18, 19^, the main pathology of angle closure disease is apposition of the peripheral iris to the trabecular meshwork impeding aqueous outflow. It is believed that dynamic movement of the iris, primarily the way the iris dilates, plays a role in angle closure pathogenesis^20–22^. Modern technology can also play its part to study the iris using computational simulations in the form of biomechanical analyses. An example would be to input ocular parameters from a patient’s optical coherence tomography (OCT) scan to evaluate how hypothetical changes in iris stiffness could influence angle closure development. These biomechanical models could evaluate the physics and predict the behaviour of the eye in any prescribed scenarios that would otherwise be difficult to analyse *in vivo*.

For example, Wang et al.^23^ explored aqueous humour dynamics to investigate its influence on the iris and pupillary block in angle closure. This was done using a physical model of the anterior chamber and tracer particles together with computational simulations. Huang et al.^24^ and Heys et al.^25^ modelled the iris-lens interactions to determine the influence of different morphological parameters (i.e. iris thickness, pupil size, anterior chamber depth, lens diameter, etc.) affecting pupillary block that would otherwise be impossible to determine experimentally. Models could also be invaluable to interpret clinical data; from dynamic OCT images, one could extract parameters such as iris stiffness and permeability^11, 12^ in addition to more traditional morphologic parameters in order to better describe the structural and biomechanical phenotype of angle closure.

However, current computational models^26–29^ of the iris are still limited in describing the physiological behaviour *in vivo*, particularly outflow of aqueous during pupil constriction and dilation. The inability to model volume changes of the iris severely limits research capabilities in angle closure, since the primary mechanism is pupillary block by the iris tissue. Our group had also previously attempted to describe the iris as a biphasic tissue; with a permeable stroma to allow for fluid exchange, and a non-linear hyperelastic model to describe both the solid phases of the stroma and the iris muscles^11, 30, 31^ However, to date, no attempts were made to isolate these tissues experimentally to describe their individual biomechanical behaviours, partly due to the difficulty in separating them *ex vivo*.

The aim of this study was to characterise the biomechanics of the main structural components of the iris (stroma and dilator muscle) experimentally. Specifically, we aimed to determine (1) the permeability of the iris stroma using a refined experimental set-up, and (2) the viscoelastic properties of both the stroma and dilator muscle through stress relaxation testing.

## Methods

For this study, we isolated two of the primary load bearing components of the iris: the stroma and the dilator muscle. The iris stroma is composed of a sponge-like meshwork^32^, porous in the middle to allow fluid movements within the anterior volume. It is also composed of fibroblasts and melanocytes that form a cellular matrix, along with vasculature, neural elements and a connective tissue extracellular matrix^33^. We isolated the stroma tissue to determine its fluid permeability, as well as the viscoelastic properties of the solid matrix.

On the other hand, the dilator muscle is made of smooth muscle cells, arranged in bundles separated by tissue septae. These interdigital systems of muscle fibres are innervated from autonomic nerve fibres within these tissue septae, communicating through other muscle fibres through large gap junctions. The surrounding cytoplasm contains a uniform distribution of myofilaments that contribute to the mechanical strength of the muscle. As the dilator muscle is impermeable due to the presence of tissue septae and gap junctions^33–35^, only its viscoelastic properties would be investigated. As the iris posterior pigmented epithelium is a component of the blood aqueous barrier, it was removed to study the stromal permeability.

### Tissue separation

Sixty freshly enucleated porcine eyes (were purchased from a local porcine abattoir (approximately aged 6 months, Singapore Food Industries Pte Ltd., Singapore). The fresh porcine eyes were refrigerated after excision, chilled in an ice box during the transport to the laboratory. Experiments were performed at room temperature of 24°C and completed within 24 hours post-mortem.

The procedure to separate the stroma and dilator tissues was similar to our previous study, which had been validated through histological analysis^31^. We made further improvements to the technique by implementing the following refinements in the procedure. First, for each eye, the entire iris was isolated from the porcine globe, and the iris pigmented epithelium gently scraped off. Next, a discontinuous cut was made to the iris. Using a mixture of tying forceps (2-500-2E, Duckworth & Kent, Hertfordshire, UK), jewellers forceps (2-900E, Duckworth & Kent, Hertfordshire, UK) and Chihara conjunctival forceps (2-500-4E, Duckworth & Kent, Hertfordshire, UK), the dilator muscle was gently lifted and peeled away from the stroma inward radially up to the sphincter, where the piece of tissue was cut off using a pair of vannas scissors (1-111, Duckworth & Kent, Hertfordshire, UK). When isolating the stroma, the dilator was removed in pieces to retain as much stroma tissue as possible, whereas when isolating the dilator muscle, care was taken to ensure no holes were present in the isolated sample, even though small amounts of stroma tissue may still be present. This method ensured proper tissue isolation to preserve the structural integrity of each sample and little to no impact on the measurement of biomechanical properties. After tissue isolation, the samples were stored in phosphate buffer saline (PBS) without calcium or magnesium to limit any potential post-mortem muscle activity^36^, diluted from 10× stock solution (17-517Q, Lonza Pharma & Biotech, Maryland, USA).

### Stroma permeability study

Isolated stroma samples (*n* = 32) underwent a flow experiment to assess their permeability. The principle of the permeability experiment was similar to that described in our previous study, whereby homogenisation of the Navier-Stokes equation^37, 38^ will yield Darcy’s law^39, 40^:

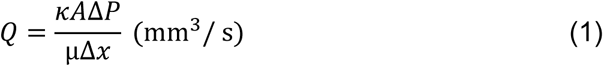

where *Q* is the flow rate (mm^3^/s), *κ* is the permeability (mm^2^), *A* is the surface area of flow (mm^2^), Δ*P* is the pressure drop across the tissue in the direction of flow (Pa), µ is the dynamic viscosity of the fluid (Pa·s) and Δ*x* is the thickness of the tissue (mm). Our measurement parameter is the hydraulic permeability, used in similar biphasic biological tissue experimentation such as the meniscus^41^. It is defined by the permeability of the tissue divided by the viscosity of the fluid (here: phosphate buffered saline or PBS):

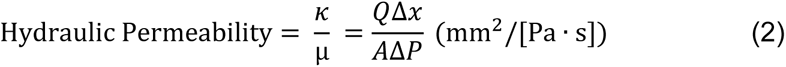

Using Darcy’s law, it was possible to obtain the hydraulic permeability of the iris stroma by setting a specific pressure difference and measuring the resulting flow rate. We designed an experimental setup consisting of a PBS column (30 mL syringe reservoir), itself connected by a tube (internal diameter (ID) = 1.00 mm) to a custom 3D-printed holder where the stroma tissue was positioned (**Figure 1**). The cross-sectional area for flow was measured using optical coherence tomography (OCT) and estimated as an ellipse (**Figure 1B and 2A**). The syringe reservoir was sufficiently large to prevent any noticeable drops in fluid level (< 1 mm) from direct flow or evaporation. The vertical height between the stroma tissue and the fluid level was approximately 204 mm to provide a pressure difference of approximately 2,000 Pa, or 15 mmHg. Outflow from the holder was through the large tube (ID = 1.00 mm), followed by a much smaller tube (ID = 0.397 mm) on the horizontal plane for the measurement of flow rate (**Figure 1C**).

**Figure 1.**
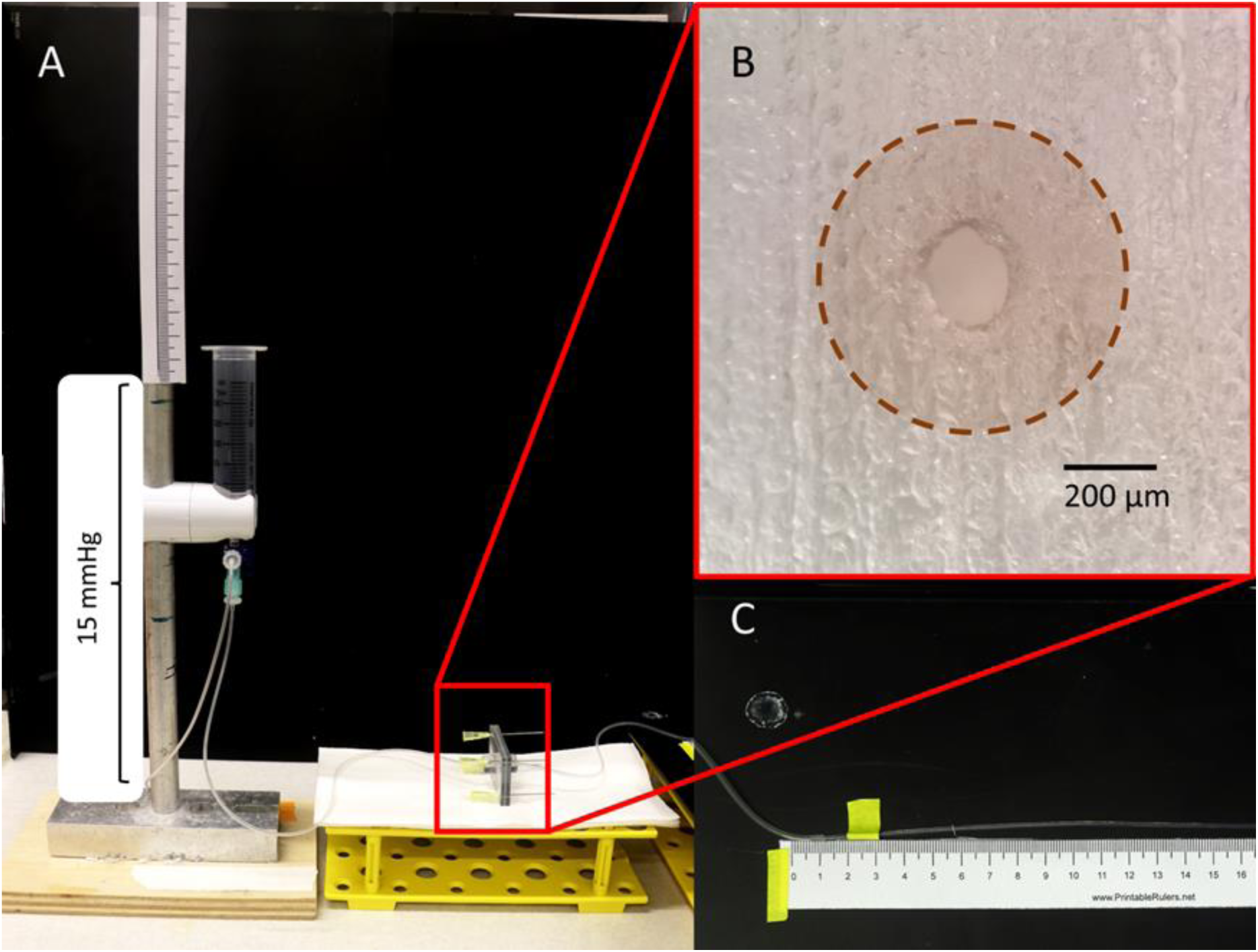
Hydraulic permeability measurement set-up. **A.** Fluid of approximately 2,000 Pa (15 mmHg) flows through a 3D printed holder. **B.** The tissue is placed between over the channel (brown dotted circle) and clamped between 2 identical 3D printed channels. The area for flow is approximately 0.0391 mm^2^, suitable for samples that are obtained from iridotomy. **C.** Flow rate through a 0.397 mm diameter tube is measured and recorded.

**Figure 2.**
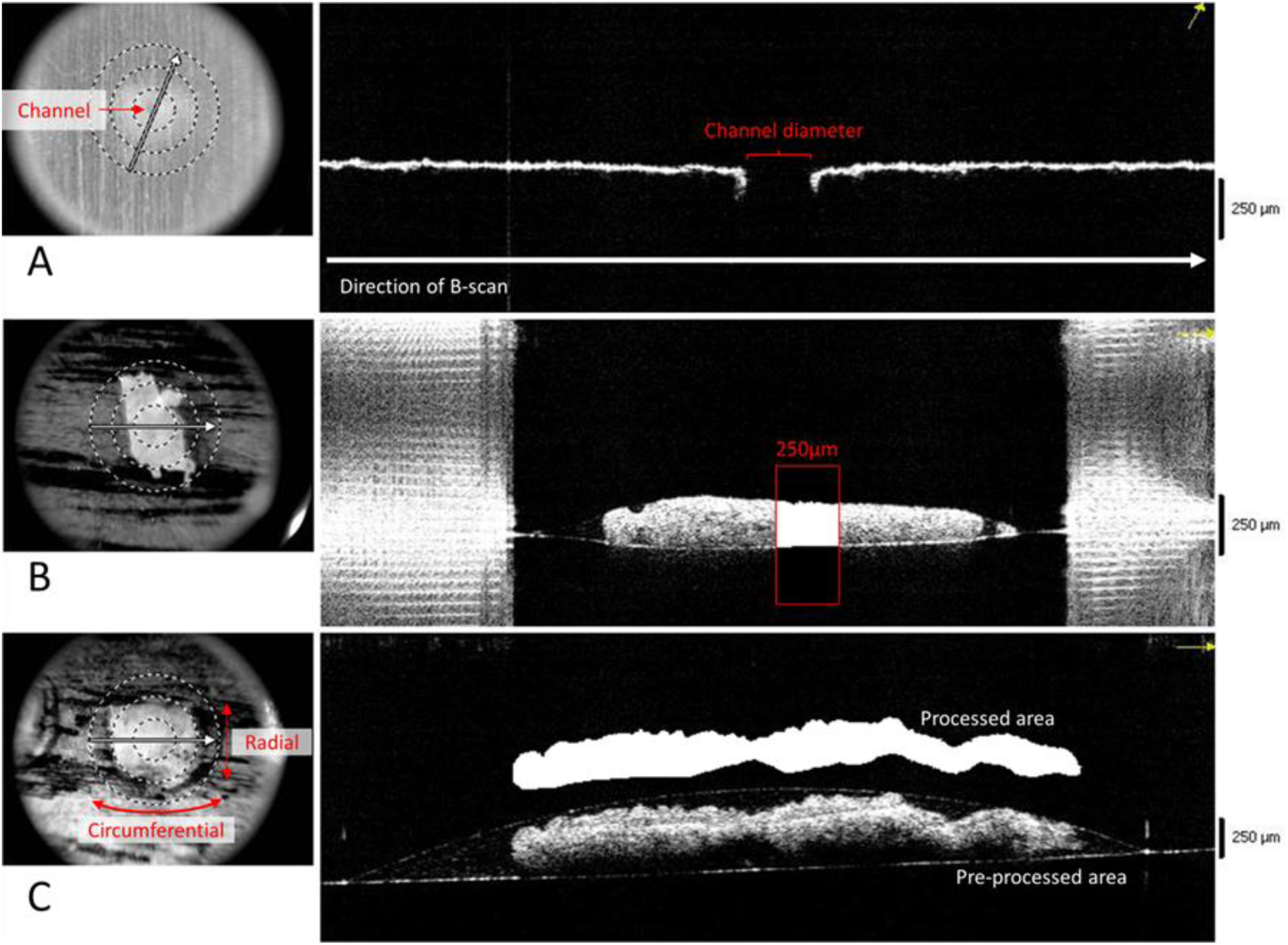
OCT images were processed to evaluate tissue parameters. The white arrows on the left depict the direction of the B-scans on the right. **A.** The channel area of the custom 3D-printed holder for the permeability experiment was estimated as an ellipse using measurements from OCT. **B.** For the permeability experiment, an area of 45 × 100 pixels was selected for each image and manually converted to binary black and white, **C.** whereas the width and area were computed for the stress relaxation experiment.

The tubes of different inner diameters were selected for the following reasons: 1) The larger tube (ID = 1.00 mm) was sufficiently large that the tube resistance was negligible; 2) The smaller tube allowed for accurate measurement of flow rate, which was measured by the time taken for 1.237 mm^3^ of fluid to flow through the stroma tissue (10 mm of the tube). A smaller tube also created outflow resistance (positive pressure), which was taken into account during the calculations. The hydraulic permeability values were reported as the mean ± standard deviation, with outliers excluded beyond 3 standard deviations.

To ensure accurate and consistent results, we calibrated the experimental setup at the start of each experiment by measuring the flow rate in the absence of tissue resistance. The measured flow rate closely matched theoretical calculations using Poiseuille’s equation, with a margin of error of less than 5%. Before testing each sample, we thoroughly flushed and inspected the entire system for bubbles and leaks to ensure the reliability of our results.

### Stress relaxation study

Iris stroma (*n* = 28) and dilator muscles (*n* = 28) underwent stress relaxation tests (i.e., stress response over time over a constant strain) using a uniaxial testing machine (5542A, Instron Corp, Norwood, MA, USA). The tested length (in the direction of the uniaxial stretch) was 1.5 mm for all stroma and dilator muscle samples, with the cross-sectional areas evaluated using OCT scans performed for each sample (**Figure 2C**) prior to the stress relaxation tests. As the iris tissues typically exhibit extremely large strains over several seconds (up to the second order of magnitude in percentage), we decided to perform the stress relaxation experiment at 50% strain. The samples were extended with a strain rate of 10% per second for 5.0 seconds and held at constant strain (ε_0_) for the next 300 seconds (**Figure 3**). All samples were clamped using sandpaper to prevent slipping (**Figure 3C**). Preconditioning was performed for 25 cycles from 0% to a maximum of 25% strain at a rate of 1% per second. Overall, samples were found to exhibit little to no hysteresis.

**Figure 3.**
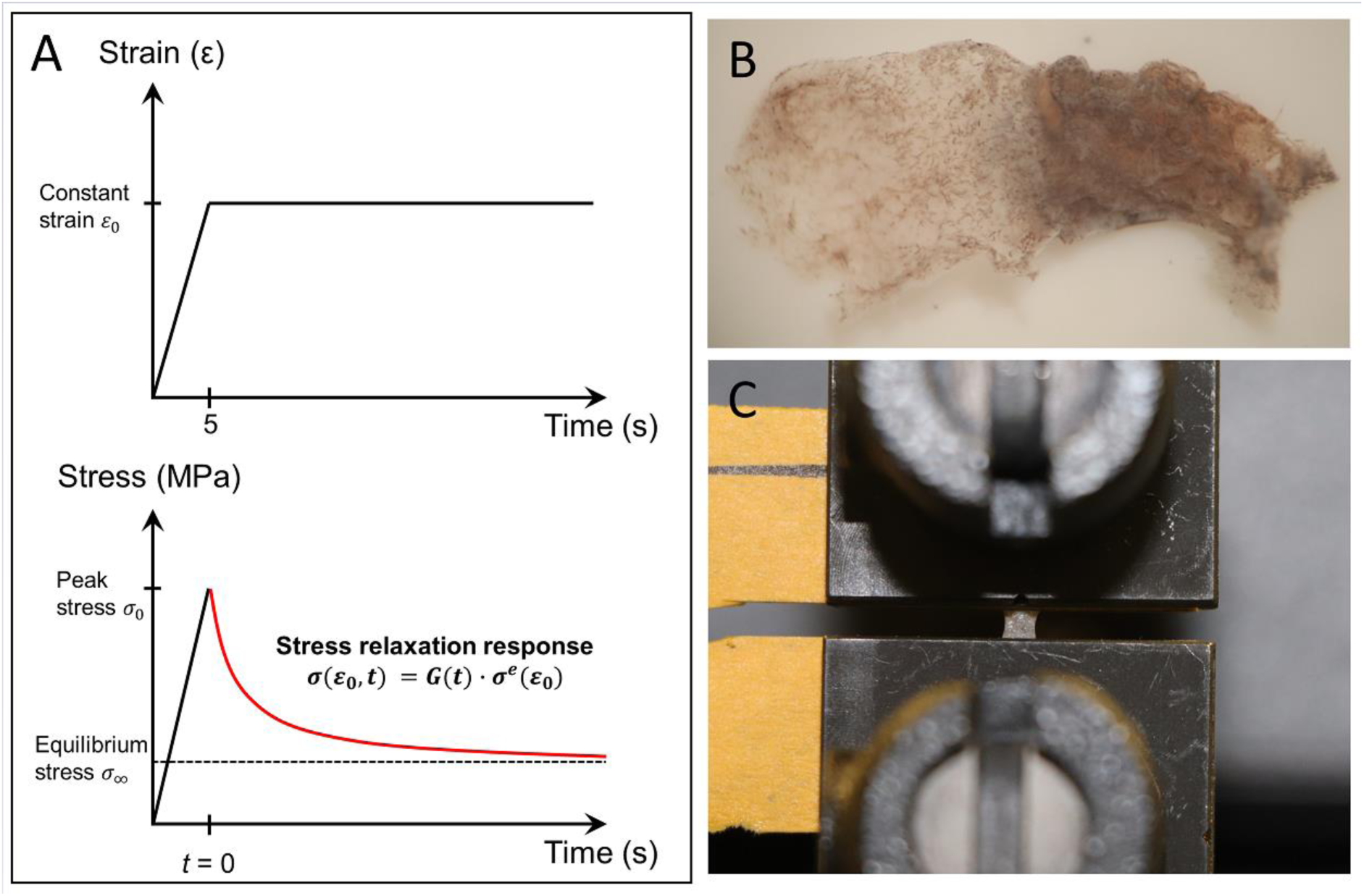
Stress relaxation experiment was conducted on the iris tissues by **A.** Applying linear strain rate of 10% per second for 5 seconds and holding constant strain as the tissue relaxes. **B.** The same protocol was performed on both the isolated dilator (left) and stroma (right) tissues. **C.** To prevent slippage, fine sandpaper was used at the clamps of the Instron machine.

To quantify the relaxation behaviour of the iris tissues, we used the linear viscoelastic theory initially described by Fung^42^ and the same mathematical model as in our previous study by Girard et al., with the stress relaxation response^43^:

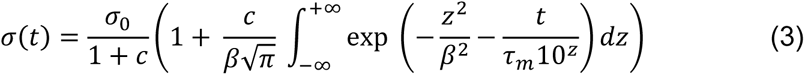

where *c* is the relaxation ratio, *β* is the half width of the Gaussian distribution^44^, *z* = log(τ/τ_*m*_), τ is the relaxation time^45^ and τ_*m*_ is the mean relaxation time constant. σ_0_ is defined as the peak stress and was directly obtained from the experimental data.

Using MATLAB (Version 2020b, Mathworks, Inc., Natick, MA, USA), we reduced noise by signal averaging and performed curve fitting of the experimental data to **Equation 3** using the differential evolution^46^ (DE) algorithm developed by Girard et al.^43^ to estimate the three model parameters: *c*, *β* and τ_*m*_. For the DE algorithm, we utilised the following parameter settings: a population size (NP) of 15, a mutation factor (F) of 0.8, a crossover probability (CR) of 0.9. The range of the variables were defined (e.g., 0 < c, β, τ_*m*_ < 10) within the DE algorithm, which represented values within the physiological range^43^. The unique global minimum determined exhibited an excellent fit, with a cost function (defined by square root of the sum of the squares) of less than 10^-3^ (**Figure 4**). The two elastic moduli, *E*_0_ and *E*_∞_ was determined by:

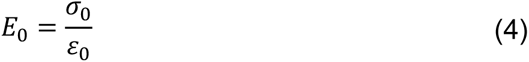

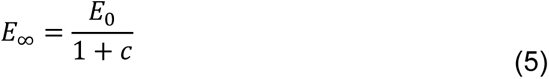

**Figure 4.**
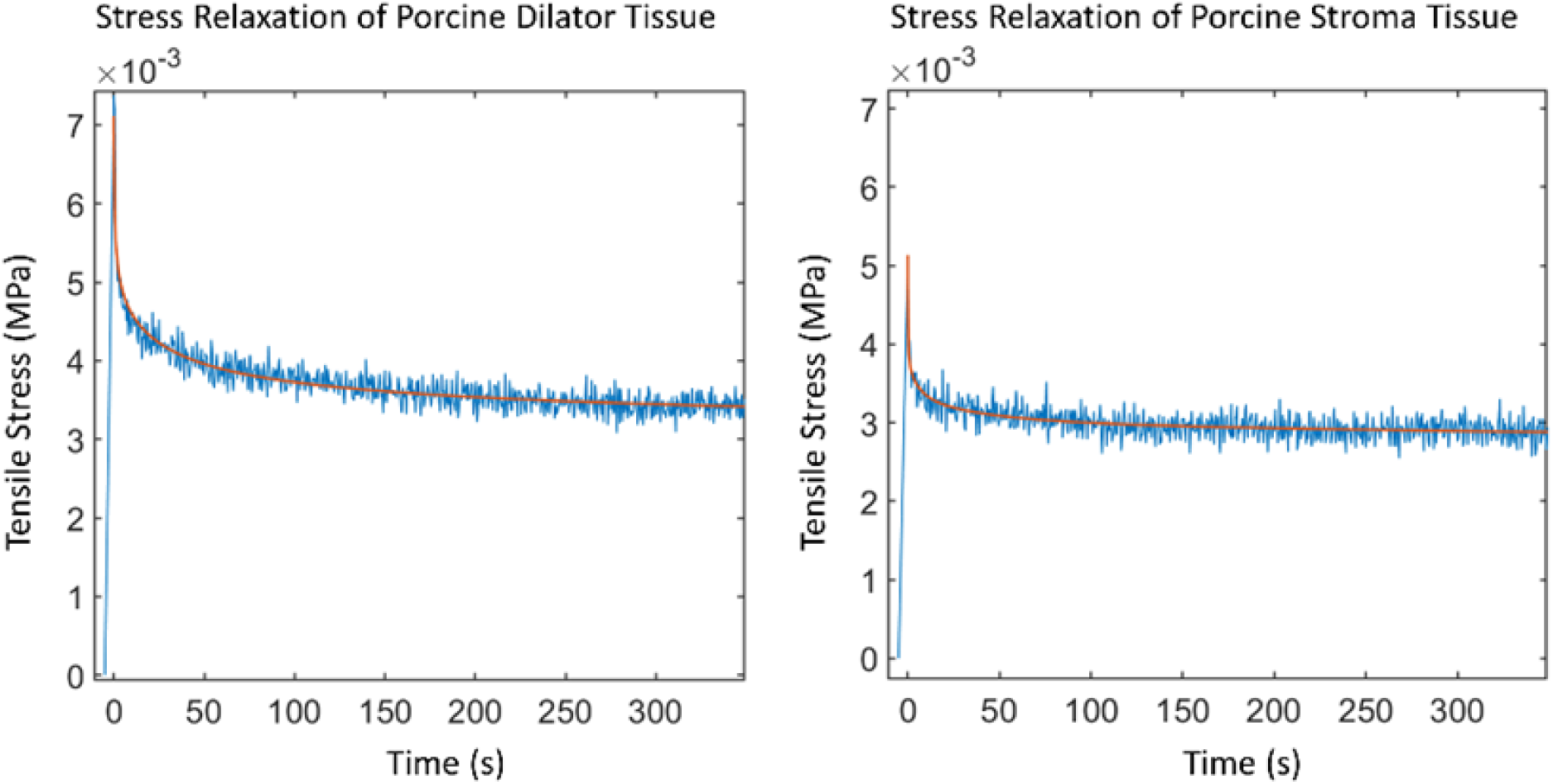
Stress relaxation behaviour of the porcine dilator tissue (left) and stroma tissue (right). The blue line represents the experimental data, and the orange line represents the fitted data.

### OCT tissue measurements

Images obtained from this study were performed using OCT (RTVue-100, Optovue Inc, Fremont, CA, USA). The stroma and dilator tissue samples were placed on a glass slide with minimal liquid to reduce influence from liquid refraction. The acquisition mode was the 8-line radial scan (each cross-sectional scan was 22.5° apart). The images were processed using Photoshop (v2022, Adobe Inc, CA, USA) and MATLAB. Thickness and cross-sectional area were evaluated using the scale from the OCT machine provided in the images (**Figure 2**).

#### Stroma permeability study

Three random images were selected per sample to determine the tissue thickness after the experiment was completed for a better reflection of the tissue state during the experiment. An area of 45 × 100 pixels was selected for each image and manually converted to binary black and white using photoshop, with the average of the 3 images computed for the tissue thickness (**Figure 3A**).

#### Stress relaxation study

Here, tissue thickness was measured prior the experiment. It was important to position the tissue uniformly so that the horizontal width of the tissue would be accurately reflected and measured by the cross-sectional images. The entire cross-section was manually converted to binary black and white using photoshop, with the width and cross-section computed for analysis (**Figure 3B**).

### Statistical Analysis

The Wilcoxon rank sum test (two-tailed) was employed to compare the five material parameters (*c*, *β*, τ_*m*_, *E*_0_ and *E*_∞_) between the dilator muscle and stroma groups (*n* = 28), to evaluate whether the tissues were inherently different.

## Results

### Stroma permeability study

The 3D tissue holder area was evaluated using OCT and determined to be 0.0391 mm^2^. The average thickness of the stroma tissue samples was 0.152 ± 0.0259 mm. The PBS flow through the stroma samples was found to be slow and laminar, with a Reynold’s number < 1, thus satisfying the Darcy’s law approximation. Specifically, flow through the experimental setup was 0.00500 ± 0.00191 mm^3^/s, exerting minimal outlet pressure (7.65 ± 2.93 Pa). The hydraulic permeability of the porcine iris stroma was then calculated as 9.49 ± 3.05 × 10^-6^ mm^2^/Pa·s, with a total of 31 samples evaluated, with 1 outlier excluded from the analysis.

### Stress relaxation study

#### Dilator muscle tissue

The average width, thickness and cross-sectional area of the dilator tissue samples were 3.78 ± 0.74 mm, 0.192 ± 0.028 mm and 0.733 ± 0.183 mm^2^ respectively. The biomechanical properties of the dilator muscle, as determined by the differential evolution algorithm, were *c* = 1.41 ± 0.60, *β* = 2.50 ± 1.40, and τ_*m*_ = 7.43 ± 4.96 s, with the two moduli calculated to be *E*_0_ = 14.14 ± 6.44 kPa and *E*_∞_ = 6.08 ± 2.74 kPa.

#### Stroma tissue

The average width, thickness and cross-sectional area of the stroma tissue samples were 3.71 ± 0.70 mm, 0.220 ± 0.04 mm and 0.830 ± 0.263 mm^2^ respectively. The biomechanical properties of the stroma, as determined by the differential evolution algorithm, were *c* = 0.707 ± 0.38, *β* = 2.06 ± 1.33, and τ_*m*_ = 1.28 ± 1.27 s, with the two moduli calculated to be *E*_0_ = 9.16 ± 3.03 kPa and *E*_∞_ = 5.54 ± 1.98 kPa.

Overall, the dilator muscle tissue exhibited greater *E*_0_ (*p* < 0.001) compared to the stroma tissue but did not show significant difference in *E*_∞_ (*p* = 0.64). The dilator muscle tissue also showed greater *c* and τ_*m*_ (*p* < 0.001) but no significant difference in *β* (*p* = 0.20).

## Discussion

In this study, we identified the stroma and dilator muscles as two structural tissues influencing iris biomechanical behaviour. To the best of our knowledge, the biomechanics of such tissue layers (when isolated) had not been studied, and we proposed herein a methodology for doing so. Overall, we found that (1) The iris stroma was highly permeable (as opposed to the dilator); (2) the instantaneous modulus (but not the equilibrium modulus) of the dilator muscle was significantly higher than that of the stroma; and that (3) the stroma exhibited a significantly shorter mean relaxation time constant as compared to the dilator muscle. Our work may ultimately help us identify an ‘healthy’ range for iris biomechanical properties in human, any deviations from which may be associated with the development of angle closure.

### The dilator muscle is stiffer than the stroma tissue in a healthy iris

In this study, we found that the instantaneous modulus of the dilator muscle was significantly higher than that of the stroma tissue – but this was not true for the equilibrium modulus. Such a difference could potentially be explained by the different fibres that constitute both tissues, including stiffer smooth muscle fibres for the dilator tissue, but a sparse collagen fibre meshwork permeable to aqueous humour for the stroma, thus allowing for greater compliance. This softer stroma layer is essential for iris accommodation, since the contraction forces (for dilation) originate from a much thinner muscle layer. Recently, it was reported that glaucomatous eyes, including eyes with primary open angle glaucoma and primary angle closure glaucoma, generally display a reduction in the pupillary light response amplitudes^47–49^. Although such alterations in the pupillary light response can be attributed to ganglion cell loss or dysfunction in the disease, we hypothesise that decreased muscle innervation and changes in biomechanical behaviour such as increased stroma stiffness, are additional underlying biological changes that may contribute to pupillometric alterations in response to light for patients with glaucoma.

### The iris accommodates movement rapidly to reach biomechanical equilibrium

In this study, we found that the mean relaxation time constants for both the dilator muscle (7.43 ± 4.96 s) and stroma (1.28 ± 1.27 s) were relatively small. These features may reflect the highly dynamic physiological properties of the iris that is constantly changing in shape in response to the accommodation reflex or the lighting environment. This is in contrast to other ocular tissues such as the corneo-scleral shell that exhibits considerably smaller changes in tissue stretch. The viscoelastic behaviour of stiffer, load bearing structural tissues (in terms of intraocular pressure) such as the sclera exhibits high moduli and greater mean relaxation time constant, whereas the small amount of muscle fibres orchestrating fine and rapid pupil dilator changes result in much smaller moduli (*E*_∞_ = 3,300 kPa for rabbit sclera vs. 5.54 kPa for iris stroma) and short mean relaxation time constants (τ_*m*_ = > 10s for rabbit sclera vs. < 10s for iris tissues)^43^.

The results also show a shorter mean relaxation time constant for the stroma tissue, with it seemingly behaving close to a linear elastic material (**Figure 4**). This could be due to its porous nature and composition; most of the stroma is composed short collagen fibrils meshwork with fibroblasts and melanosomes^33, 34^. The stroma matrix deforms rapidly in both extension and compression as aqueous humour gets quickly displaced through its pores, reaching structural equilibrium within seconds. Conversely, the dilator muscle takes longer to reach biomechanical equilibrium, the simple bundles of smooth muscle cells^33^ containing actin and myosin filaments passively extend irreversibly, until the filaments are activated to overlap and contract during pupil constriction.

### Changes in iris tissue properties can affect iris curvature and induce angle closure

In angle closure, the behaviour and shape of the iris in the mydriatic state affects the chamber angle. Patients with angle closure glaucoma were found to have anterior convex-to-convex (in dark and light) iris configuration^50^, possibly signalling a biomechanical mismatch between the iris stroma and muscle tissues. There are several potential mechanisms that can result in an anteriorly convex iris. Firstly, if the iris stroma undergoes stiffening, the tissue may bow upward to counteract compression by the dilator muscle. Secondly, changes in the composition of the iris stroma could lead to an increased matrix-to-aqueous ratio, reducing compressibility and resulting in anterior bowing. Thirdly, a decrease in stromal permeability could impede the outflow of aqueous humour from the iris^32^, leading to volume retention^51^ and increased iris thickness^52^, ultimately causing convexity. This change would likely be attributed to morphological changes at the anterior boundary layer, where the density of the tissue is the primary limiting factor restricting aqueous fluid migration. Finally, a mismatch in biomechanical behaviour between the stroma and surrounding muscles, such as an increase in stroma mean relaxation time constant, could affect how the iris behaves in unison. In the case of pupil dilation, an anteriorly convex iris could result initially before the stresses are evenly distributed, only becoming less convex over a longer period of time.

The lack of iris features such as crypts and furrows^53^ had been linked to the development of angle closure. The presence of crypts, which are diamond shape lacunae causing discontinuities on the anterior boundary layer, could facilitate aqueous humour movement across the stroma tissue and prevent the iris from bulging and bowing during pupil dilation. The presence of lower number of crypts had been found to be proportional to the incidence of angle closure in patients^54^. Furrows, which are circumferential creases where the iris folds at the periphery during pupil dilation, could also be associated with the prevalence of angle closure. With longer furrow lengths, it was found that the irides were more likely to have greater volume and less aqueous loss during pupil dilation^55^. These iris features provide not only greater insight into their associations with angle closure, but also possible reasons for the deviations from mean iris behaviour that could be quantified biomechanically.

### Our work allows us to better describe the passive extensive behaviour of the ex vivo porcine iris

This study represented the iris as a structural bilayer; the anterior stroma as a solid-liquid biphasic tissue with solid viscoelastic properties and fluid permeability, and the posterior dilator muscle as a solid with viscoelastic properties. We believe this captures the unique behaviour of the iris, which is unlike many other tissues in the body. The iris experiences large deformations created by the posteriorly positioned muscles, coupled with interaction with the surrounding fluid medium from the partially porous and anteriorly positioned stroma. By isolating the individual primary load bearing layers, it was possible to accurately determine their biomechanical behaviour. Previous research into the iris took different approaches: Whitcomb et al. performed nano-indentation of the porcine iris anterior and posterior segments to determine overall iris stiffness^28^, while Narayanaswamy et al. used atomic force microscopy^56^. There are some anatomical and biomechanical points to consider for these approaches. First, for the anterior indentation, the small level of indentation in a porous stroma could effectively be flexing fibres between the gaps of the tissue, undermining the true tissue stiffness. Second, when the antagonistic muscles contract, the iris is either in a compressive or extensive state in the radial direction, orthogonal to the direction of indentation. The assumption of an isotropic tissue may not be biomechanically accurate to represent the radially oriented dilator muscles, and it may be critical to test the tissues according to their preferred fibre orientations as we had performed herein.

Whilst the method for measuring biomechanical properties is only possible *ex vivo*, we believe that capturing a better representation of the iris behaviour is invaluable for clinical research. First, we could utilise this technique to investigate the tissue biomechanical differences between normal and angle closure iridectomy samples. Next, through evaluation of miosis and mydriasis OCT data, we believe that iris biomechanical differences in angle closure can be validated *in vivo* using computational methods such as inverse finite element^11^ and AI-driven approaches to access tissue biomechanics. From a clinical perspective, our goal is to identify differences in biomechanical data between normal and disease subjects by evaluating complex OCT data, and the understanding of differences can be combined with AI to improve disease prognosis.

### Limitations

In this study, several limitations warrant further discussion. First, a more complete biomechanical breakdown of the iris tissue properties should also include the sphincter muscle. The omission of the sphincter muscle was due to the difficulty in isolating this circular tissue. However, it is theoretically possible to equate the passive biomechanical behaviour of both the dilator and sphincter muscle since they are antagonistic tissues. The differences in speeds of pupil constriction and dilation suggest possible small tissue differences, perhaps due to different nerve innervations, but microscopic studies have shown that these muscles are typical smooth muscles^33^, and hence are likely to exhibit similar biomechanical properties. Moreover, a better approach would involve incorporating the biphasic poroviscoelastic (BPVE) constitutive model, which accounts for both viscoelasticity and permeability. In our study, we focused on evaluating the solid linear viscoelastic theory for modelling the iris dilator muscle and the iris stroma. Additionally, we separately assessed the hydraulic permeability of the iris stroma due to our previous study’s^31^ findings suggesting that the high permeability had little impact on tissue behaviour (Appendix A). This corroborates the results investigated by Greiner et al.^57^, who found that the fluid nominal stresses were negligible for permeability values above 10^-6^ mm^2^/Pa·s. However, it is advisable for future investigations to consider employing the solid linear viscoelastic theory for the muscle tissue and the BPVE theory for the stroma tissue, especially when considering other loading scenarios.

Second, a limitation of experimental *ex vivo* biomechanical determination of tissues is the inability to determine active muscle properties. Changes in pupil consists of at least three tissue changes; pupil constriction requires active sphincter muscle contraction, dilator muscle extension (relaxation) and stroma extension, whilst pupil dilation requires active dilator muscle contraction, sphincter muscle extension (relaxation) and stroma compression. Whilst it would be a logistical challenge to determine active muscle forces by keeping the tissues alive, such forces might be able to be estimated with *in vivo* imaging such as OCT.

Third, the stroma undergoes both compressive and tensile stresses with pupillary changes, however, on our study we only performed tensile experiments. In addition, permeability could be both affected by tensile and compressive stress, as is true for other soft tissues such as cartilage^58, 59^. Novel mechanical test equipment might need to be designed to assess such properties.

Fourth, we had loaded the tissue at a rate of 10% per second for 5 seconds. Our original tissue length measured approximately 4 – 4.5 mm, trimmed to 1.5 mm. Hence our loading rate corresponds to a constriction rate of approximately 0.4 – 0.45 mm/s. *In vivo* constriction rates^60, 61^ range can be as slow as 0.5 mm/s and increase up to 7.5 mm/s depending on light intensity, with the maximum rate achieved at 550 cd/m^2^. We modelled our loading rate based on physiological dynamic accommodation constriction rates^62^, which also corresponded to the lower limits of dark-to-light constriction rates. It would be important for future studies to investigate the upper limits of constriction rates for a more comprehensive analysis of the iris’ biomechanical behaviour.

## Conclusion

Determining the biomechanical properties of the individual iris layers may be invaluable in uncovering how and why the iris deforms in the anterior chamber in normal and eyes with angle closure. This porcine study established a protocol to be replicated on human iris tissues, by isolating the individual components for analysis. In learning more about iris biomechanics, we hope to establish a better understanding of the pathogenesis of angle closure glaucoma.

## Acknowledgments

The authors thank the donors of the National Glaucoma Research, a program of the BrightFocus Foundation, for support of this research (G2021010S [MJAG]); the NMRC-LCG grant ‘TAckling & Reducing Glaucoma Blindness with Emerging Technologies (TARGET)’, award ID: MOH-OFLCG21jun-0003 [MJAG]; and the “Retinal Analytics through Machine learning aiding Physics (RAMP)" project that is supported by the National Research Foundation (NRF2019-THE002-0006 [MJAG]).

---

**Table 1.**
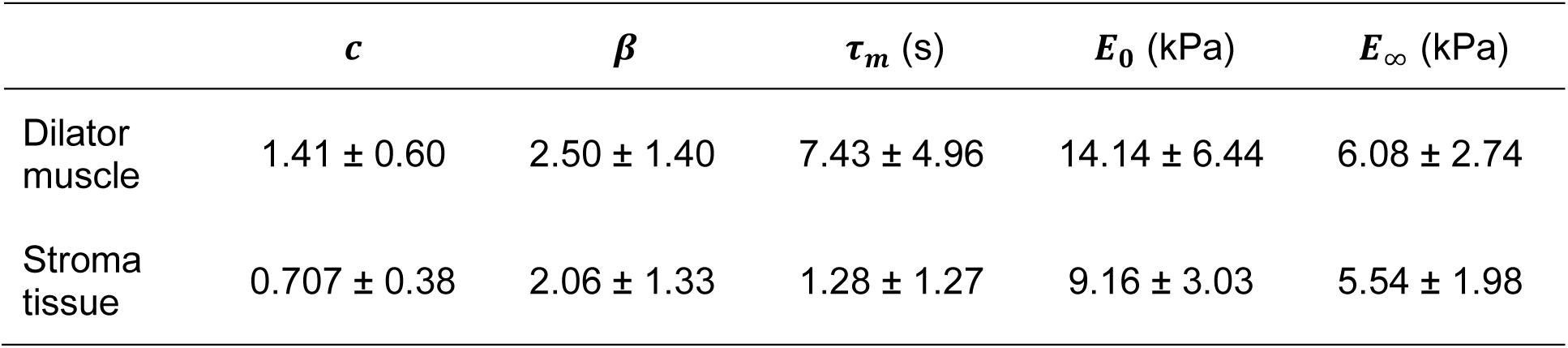
Viscoelastic parameters of the iris stroma and dilator muscle tissues (mean value ± standard deviation).

## Notes

### Competing Interest Statement

The authors have declared no competing interest.

### Summary of Updates

Updated to include comments from reviewers

