## Appendix A for "The Structural Layers of the Porcine Iris Exhibit Inherently Different Biomechanical Properties"

The biphasic poroviscoelastic theory (BPVE) has been extensively used to model the stress-strain behaviour of soft tissues, governed by the continuity equation:

$$\mathrm{div} \cdot (\mathrm{v}^s + \mathrm{w}) = 0$$

whereby $\mathrm{v}^s$ is the solid matrix velocity and $\mathrm{w}$ is the fluid flux relative to the solid. At high permeability values, fluid leaves the tissue ($\mathrm{w}$ falls rapidly to 0) (refer to Figure F below) almost instantaneously, hence the contribution of the stress relaxation response of the tissue can be approximated to the viscoelastic effect of the solid phase only. The fluid flux is determined by Darcy's law: $\mathrm{w} = -\mathrm{k} \cdot (\mathrm{grad}\, p - \rho_T^w \mathrm{b}^w)$, where $p$ is the fluid pressure, $\rho_T^w$ is the fluid density and $\mathrm{b}^w$ is the external body force per mass acting on the fluid (zero during stress relaxation).

The total stress in the mixture is $\sigma = -pI + \sigma^e$, where $I$ is the identity tensor and $\sigma^e$ is the stress of the solid, which is represented by the viscoelastic model below. The mixture momentum balance is $\mathrm{div}\, \sigma = 0$, in the absence of body forces.

To show the potential influence of permeability on our stress relaxation experiments, we performed a computational study using the biphasic model in FEBio (FEBio Studio v1.8.0, University of Utah, UT, USA) using the governing equations above.

The finite element model was a tissue measuring 1.5 (L) x 3.0 (W) x 0.2 (T) mm similar to the stroma samples isolated in the experiment. The material model consisted of the following parameters and definitions:

Solid volume fraction: 0.6
Viscoelastic coefficients (from box spectrum model with n=1 for simplicity):
c = 2.8, $\tau$ = 1.28s
Isotropic elastic solid modulus: 1.9 kPa
Poisson's ratio: 0.3
Permeability: $10^{-4}$, $10^{-5}$, $10^{-8}$, $10^{-10}$, $10^{-11}$, $10^{-14}$ $mm^2$/Pa.s

The computational analysis consisted of the following definitions: uniaxial extension at a rate of 10% per second for 5 seconds, followed by a stress relaxation duration of 100 seconds. The analysis was conducted using a transient analysis type with a solver employing a non-symmetric matrix and the Broyden solver.

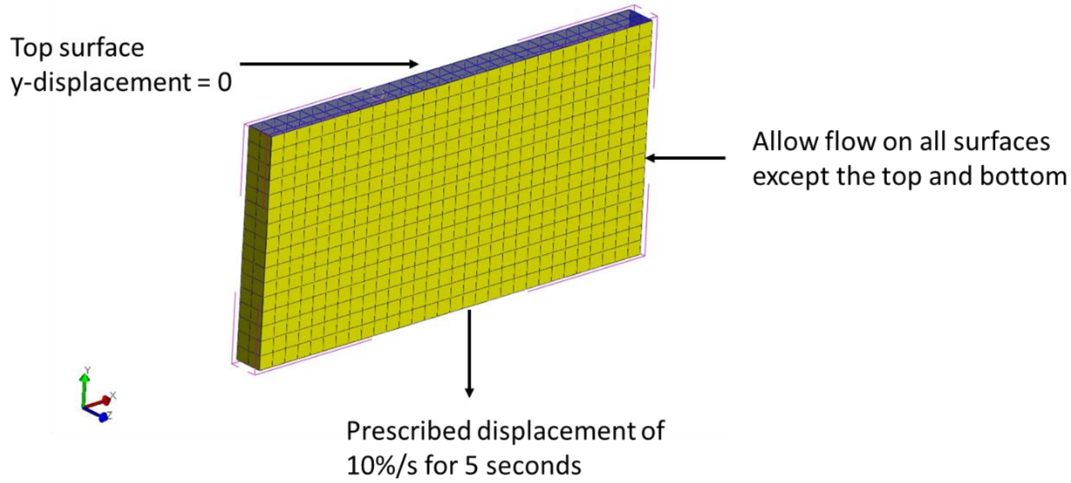


Figure A. Boundary conditions for the finite element model.

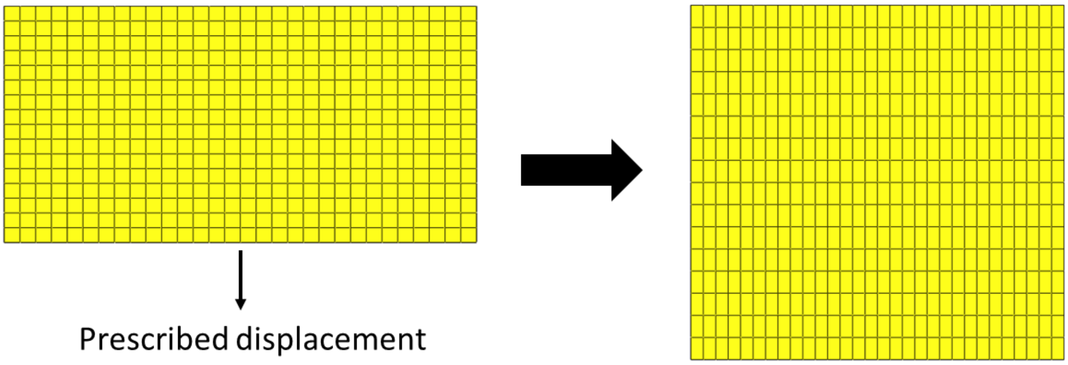


Figure B. Before (left) and after (right) prescribed displacement of the model at the strain rate of 10% per second for 5 seconds.

Our results showed that at permeability values higher than $10^{-8}$ $mm^2/Pa.s$, fluid flow (out of the stroma) was rapid, contributing to less than 0.22% of the total stresses within the tissue. This suggests that the stress-strain behaviour of the stroma could be approximated solely by the solid phase of the biphasic tissue.

Since our experimental permeability values (Figure C and D) for the iris stroma were on the order of $10^{-5}$ $mm^2/Pa.s$ (and thus higher than $10^{-8}$ $mm^2/Pa.s$), it would be safe to say that BPVE is not necessary for our chosen loading conditions. In fact, our simulations suggested that fluid flow only accounted for a negligible fraction ($6x10^{-8}$%) of the total stresses for such permeability values (i.e. $10^{-5}$ $mm^2/Pa.s$).

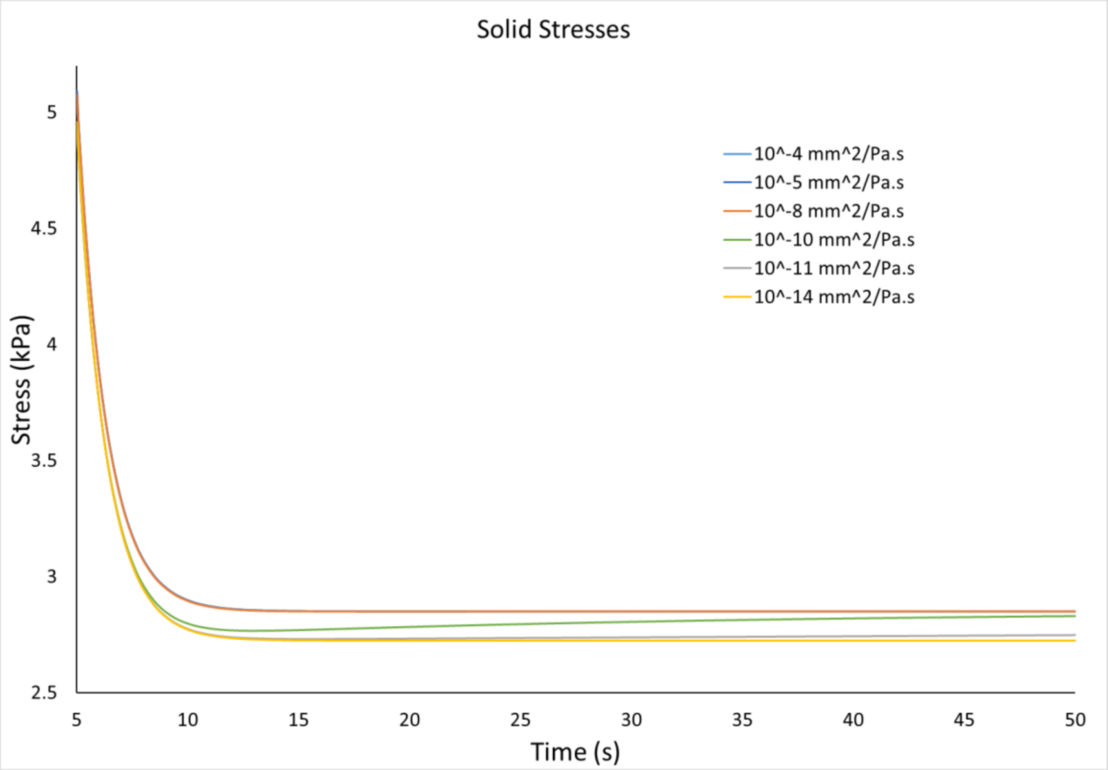


Figure C. Stresses within the solid phase of the simulated stroma tissue model.

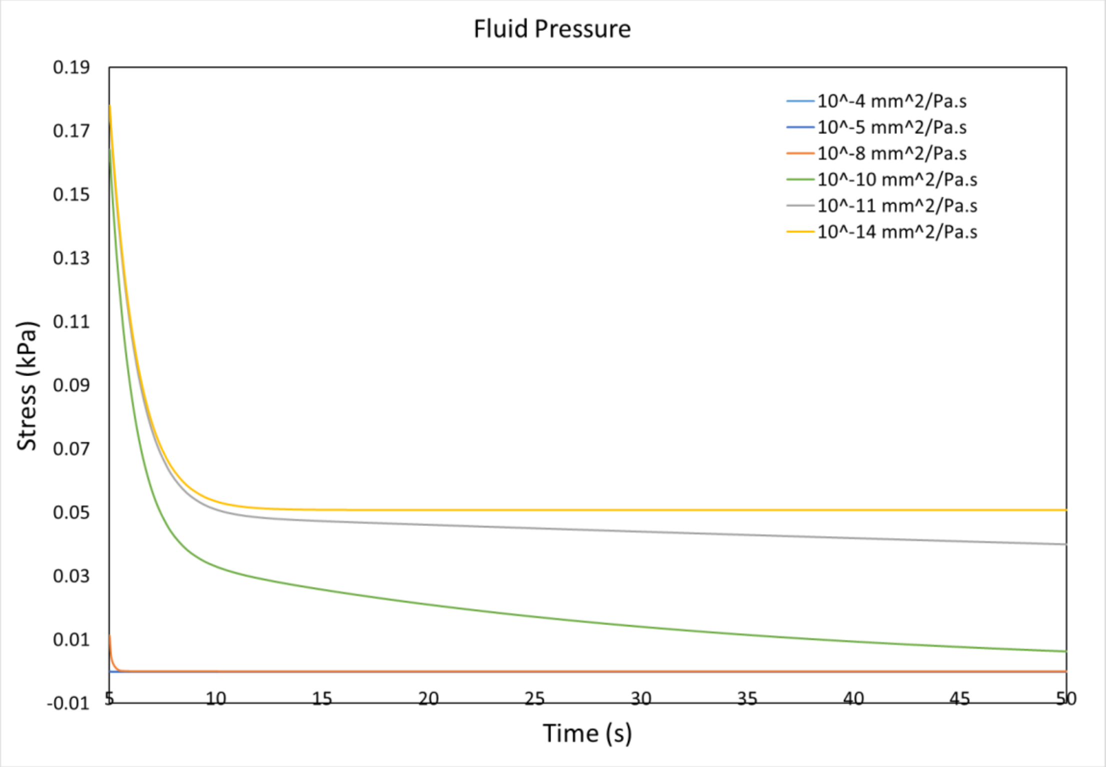


Figure D. Stresses within the fluid phase of the simulated stroma tissue model.

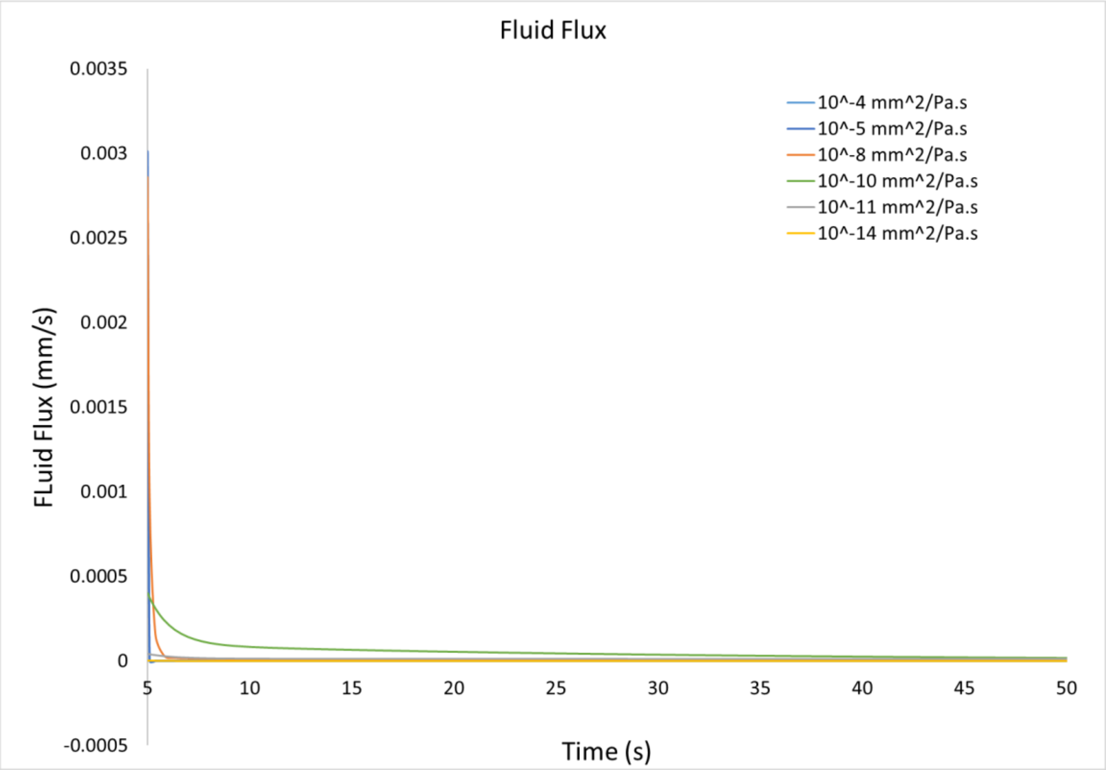


Figure E. Flux of the fluid along the boundary where flow was allowed.

Raw Data (After 5s ramp to 50s)

| Time | Solid Stresses (kPa) 10$^{-4}$ mm$^2$/Pa.s | 10$^{-5}$ mm$^2$/Pa.s | 10$^{-8}$ mm$^2$/Pa.s | 10$^{-10}$ mm$^2$/Pa.s | 10$^{-11}$ mm$^2$/Pa.s | 10$^{-14}$ mm$^2$/Pa.s |
|---|---|---|---|---|---|---|
| 5 | 5.09303 | 5.09301 | 5.07609 | 4.9404 | 4.95428 | 4.95702 |
| 5.1 | 4.92696 | 4.92695 | 4.91452 | 4.77793 | 4.78929 | 4.79168 |
| 5.38 | 4.52451 | 4.5245 | 4.51475 | 4.38345 | 4.38939 | 4.39101 |
| 5.804 | 4.05848 | 4.05848 | 4.05051 | 3.92597 | 3.92626 | 3.92703 |
| 6.3432 | 3.64819 | 3.64819 | 3.64175 | 3.52324 | 3.51853 | 3.51855 |
| 6.97456 | 3.34112 | 3.34112 | 3.33591 | 3.2226 | 3.21348 | 3.21283 |
| 7.67965 | 3.13552 | 3.13552 | 3.13126 | 3.02252 | 3.00939 | 3.00814 |
| 8.44372 | 3.00863 | 3.00863 | 3.00511 | 2.90045 | 2.88364 | 2.8818 |
| 9.25497 | 2.93499 | 2.93499 | 2.93206 | 2.83113 | 2.81087 | 2.80849 |
| 10.104 | 2.89423 | 2.89423 | 2.89178 | 2.79433 | 2.77084 | 2.76791 |
| 10.9832 | 2.87249 | 2.87249 | 2.87044 | 2.77629 | 2.74972 | 2.74627 |
| 11.8865 | 2.86123 | 2.86122 | 2.85951 | 2.76855 | 2.73904 | 2.73505 |
| 12.8092 | 2.85552 | 2.85552 | 2.85409 | 2.76623 | 2.73388 | 2.72937 |
| 13.7474 | 2.85268 | 2.85268 | 2.85149 | 2.76667 | 2.73159 | 2.72655 |
| 14.6979 | 2.85129 | 2.85129 | 2.8503 | 2.76845 | 2.73074 | 2.72516 |
| 15.6583 | 2.85062 | 2.85062 | 2.8498 | 2.77086 | 2.7306 | 2.72449 |
| 16.6267 | 2.85029 | 2.85029 | 2.84961 | 2.77352 | 2.73081 | 2.72417 |
| 17.6013 | 2.85014 | 2.85014 | 2.84958 | 2.77626 | 2.73119 | 2.72402 |
| 18.5811 | 2.85006 | 2.85006 | 2.8496 | 2.77899 | 2.73165 | 2.72395 |
| 19.5649 | 2.85003 | 2.85003 | 2.84965 | 2.78166 | 2.73216 | 2.72391 |
| 20.5519 | 2.85001 | 2.85001 | 2.8497 | 2.78425 | 2.73267 | 2.7239 |
| 21.5415 | 2.85001 | 2.85001 | 2.84974 | 2.78677 | 2.7332 | 2.72389 |
| 22.5332 | 2.85 | 2.85 | 2.84979 | 2.78919 | 2.73373 | 2.72389 |
| 23.5266 | 2.85 | 2.85 | 2.84982 | 2.79154 | 2.73426 | 2.72389 |
| 24.5212 | 2.85 | 2.85 | 2.84985 | 2.79379 | 2.73479 | 2.72389 |
| 25.517 | 2.85 | 2.85 | 2.84988 | 2.79597 | 2.73532 | 2.72389 |
| 26.5136 | 2.85 | 2.85 | 2.8499 | 2.79806 | 2.73584 | 2.72389 |
| 27.5109 | 2.85 | 2.85 | 2.84992 | 2.80007 | 2.73637 | 2.72389 |
| 28.5087 | 2.85 | 2.85 | 2.84993 | 2.80201 | 2.73689 | 2.72389 |
| 29.507 | 2.85 | 2.85 | 2.84994 | 2.80388 | 2.73741 | 2.72389 |
| 30.5056 | 2.85 | 2.85 | 2.84995 | 2.80567 | 2.73792 | 2.72389 |
| 31.5045 | 2.85 | 2.85 | 2.84996 | 2.8074 | 2.73844 | 2.72389 |
| 32.5036 | 2.85 | 2.85 | 2.84997 | 2.80906 | 2.73895 | 2.72389 |
| 33.5029 | 2.85 | 2.85 | 2.84997 | 2.81065 | 2.73946 | 2.72389 |
| 34.5023 | 2.85 | 2.85 | 2.84998 | 2.81219 | 2.73997 | 2.72389 |
| 35.5018 | 2.85 | 2.85 | 2.84998 | 2.81366 | 2.74048 | 2.72389 |
| 36.5015 | 2.85 | 2.85 | 2.84999 | 2.81508 | 2.74098 | 2.72389 |
| 37.5012 | 2.85 | 2.85 | 2.84999 | 2.81645 | 2.74149 | 2.72389 |
| 38.5009 | 2.85 | 2.85 | 2.84999 | 2.81776 | 2.74199 | 2.72389 |
| 39.5007 | 2.85 | 2.85 | 2.84999 | 2.81902 | 2.74248 | 2.72389 |

| | | | | | | |
|---|---|---|---|---|---|---|
| 40.5006 | 2.85 | 2.85 | 2.84999 | 2.82023 | 2.74298 | 2.72389 |
| 41.5005 | 2.85 | 2.85 | 2.84999 | 2.8214 | 2.74347 | 2.7239 |
| 42.5004 | 2.85 | 2.85 | 2.85 | 2.82252 | 2.74396 | 2.7239 |
| 43.5003 | 2.85 | 2.85 | 2.85 | 2.8236 | 2.74445 | 2.7239 |
| 44.5002 | 2.85 | 2.85 | 2.85 | 2.82463 | 2.74494 | 2.7239 |
| 45.5002 | 2.85 | 2.85 | 2.85 | 2.82563 | 2.74542 | 2.7239 |
| 46.5002 | 2.85 | 2.85 | 2.85 | 2.82658 | 2.74591 | 2.7239 |
| 47.5001 | 2.85 | 2.85 | 2.85 | 2.8275 | 2.74639 | 2.7239 |
| 48.5001 | 2.85 | 2.85 | 2.85 | 2.82838 | 2.74687 | 2.7239 |
| 49.5001 | 2.85 | 2.85 | 2.85 | 2.82923 | 2.74734 | 2.7239 |
| 50.5001 | 2.85 | 2.85 | 2.85 | 2.83005 | 2.74782 | 2.7239 |

| | Fluid Stresses (kPa) | | | | | |
|---|---|---|---|---|---|---|
| Time | $10^{-4}$ $mm^2$/Pa.s | $10^{-5}$ $mm^2$/Pa.s | $10^{-8}$ $mm^2$/Pa.s | $10^{-10}$ $mm^2$/Pa.s | $10^{-11}$ $mm^2$/Pa.s | $10^{-14}$ $mm^2$/Pa.s |
| 5 | 1.19E-06 | 1.19E-05 | 0.011289 | 0.164117 | 0.176711 | 0.17778 |
| 5.1 | 6.30E-11 | 6.29E-09 | 0.003852 | 0.152998 | 0.167103 | 0.168346 |
| 5.38 | 6.85E-13 | 6.95E-11 | 0.000623 | 0.127022 | 0.143898 | 0.145493 |
| 5.804 | 6.36E-13 | 6.36E-11 | 0.000118 | 0.098333 | 0.117146 | 0.119042 |
| 6.3432 | 5.77E-13 | 5.77E-11 | 6.17E-05 | 0.074353 | 0.093701 | 0.095765 |
| 6.97456 | 5.12E-13 | 5.12E-11 | 5.16E-05 | 0.057272 | 0.076223 | 0.07835 |
| 7.67965 | 4.47E-13 | 4.47E-11 | 4.50E-05 | 0.046245 | 0.064543 | 0.066692 |
| 8.44372 | 3.86E-13 | 3.85E-11 | 3.90E-05 | 0.039483 | 0.057314 | 0.059495 |
| 9.25497 | 3.29E-13 | 3.29E-11 | 3.34E-05 | 0.035354 | 0.053068 | 0.055312 |
| 10.104 | 2.78E-13 | 2.78E-11 | 2.84E-05 | 0.032714 | 0.050645 | 0.052989 |
| 10.9832 | 2.34E-13 | 2.34E-11 | 2.40E-05 | 0.030866 | 0.049266 | 0.051741 |
| 11.8865 | 1.95E-13 | 1.95E-11 | 2.02E-05 | 0.029426 | 0.048454 | 0.051084 |
| 12.8092 | 1.63E-13 | 1.63E-11 | 1.69E-05 | 0.028197 | 0.04794 | 0.05074 |
| 13.7474 | 1.35E-13 | 1.35E-11 | 1.41E-05 | 0.027079 | 0.047578 | 0.050557 |
| 14.6979 | 1.12E-13 | 1.12E-11 | 1.17E-05 | 0.02603 | 0.04729 | 0.050455 |
| 15.6583 | 9.21E-14 | 9.20E-12 | 9.74E-06 | 0.025027 | 0.047039 | 0.050394 |
| 16.6267 | 7.58E-14 | 7.58E-12 | 8.08E-06 | 0.024063 | 0.046806 | 0.050352 |
| 17.6013 | 6.24E-14 | 6.24E-12 | 6.69E-06 | 0.023133 | 0.046581 | 0.050319 |
| 18.5811 | 5.13E-14 | 5.13E-12 | 5.53E-06 | 0.022235 | 0.04636 | 0.050291 |
| 19.5649 | 4.21E-14 | 4.21E-12 | 4.57E-06 | 0.02137 | 0.046141 | 0.050265 |
| 20.5519 | 3.46E-14 | 3.46E-12 | 3.78E-06 | 0.020536 | 0.045923 | 0.05024 |
| 21.5415 | 2.84E-14 | 2.83E-12 | 3.12E-06 | 0.019732 | 0.045706 | 0.050215 |
| 22.5332 | 2.32E-14 | 2.32E-12 | 2.57E-06 | 0.018959 | 0.04549 | 0.050191 |
| 23.5266 | 1.90E-14 | 1.90E-12 | 2.12E-06 | 0.018214 | 0.045275 | 0.050167 |
| 24.5212 | 1.56E-14 | 1.56E-12 | 1.75E-06 | 0.017498 | 0.04506 | 0.050142 |
| 25.517 | 1.28E-14 | 1.28E-12 | 1.44E-06 | 0.016809 | 0.044846 | 0.050118 |
| 26.5136 | 1.05E-14 | 1.05E-12 | 1.19E-06 | 0.016147 | 0.044633 | 0.050094 |
| 27.5109 | 8.57E-15 | 8.57E-13 | 9.80E-07 | 0.015511 | 0.044421 | 0.050069 |
| 28.5087 | 7.02E-15 | 7.01E-13 | 8.07E-07 | 0.014899 | 0.04421 | 0.050045 |

| | | | | | | |
|---|---|---|---|---|---|---|
| 29.507 | 5.74E-15 | 5.74E-13 | 6.65E-07 | 0.014311 | 0.044 | 0.050021 |
| 30.5056 | 4.70E-15 | 4.70E-13 | 5.48E-07 | 0.013746 | 0.043791 | 0.049996 |
| 31.5045 | 3.85E-15 | 3.85E-13 | 4.52E-07 | 0.013204 | 0.043582 | 0.049972 |
| 32.5036 | 3.15E-15 | 3.15E-13 | 3.72E-07 | 0.012682 | 0.043375 | 0.049948 |
| 33.5029 | 2.58E-15 | 2.58E-13 | 3.06E-07 | 0.012181 | 0.043168 | 0.049923 |
| 34.5023 | 2.11E-15 | 2.11E-13 | 2.52E-07 | 0.0117 | 0.042963 | 0.049899 |
| 35.5018 | 1.73E-15 | 1.73E-13 | 2.08E-07 | 0.011238 | 0.042758 | 0.049875 |
| 36.5015 | 1.41E-15 | 1.41E-13 | 1.71E-07 | 0.010794 | 0.042555 | 0.049851 |
| 37.5012 | 1.16E-15 | 1.16E-13 | 1.41E-07 | 0.010367 | 0.042352 | 0.049826 |
| 38.5009 | 9.46E-16 | 9.46E-14 | 1.16E-07 | 0.009958 | 0.04215 | 0.049802 |
| 39.5007 | 7.74E-16 | 7.74E-14 | 9.57E-08 | 0.009564 | 0.041949 | 0.049778 |
| 40.5006 | 6.33E-16 | 6.33E-14 | 7.88E-08 | 0.009186 | 0.04175 | 0.049754 |
| 41.5005 | 5.18E-16 | 5.18E-14 | 6.49E-08 | 0.008823 | 0.041551 | 0.04973 |
| 42.5004 | 4.24E-16 | 4.24E-14 | 5.35E-08 | 0.008474 | 0.041353 | 0.049705 |
| 43.5003 | 3.47E-16 | 3.47E-14 | 4.41E-08 | 0.008139 | 0.041156 | 0.049681 |
| 44.5002 | 2.84E-16 | 2.84E-14 | 3.63E-08 | 0.007818 | 0.04096 | 0.049657 |
| 45.5002 | 2.32E-16 | 2.32E-14 | 2.99E-08 | 0.007509 | 0.040765 | 0.049633 |
| 46.5002 | 1.90E-16 | 1.90E-14 | 2.46E-08 | 0.007212 | 0.04057 | 0.049609 |
| 47.5001 | 1.56E-16 | 1.56E-14 | 2.03E-08 | 0.006927 | 0.040377 | 0.049585 |
| 48.5001 | 1.27E-16 | 1.27E-14 | 1.67E-08 | 0.006653 | 0.040185 | 0.04956 |
| 49.5001 | 1.04E-16 | 1.04E-14 | 1.38E-08 | 0.00639 | 0.039993 | 0.049536 |
| 50.5001 | 8.53E-17 | 8.53E-15 | 1.13E-08 | 0.006138 | 0.039803 | 0.049512 |

| | Fluid Flux (mm/s) | | | | | |
|---|---|---|---|---|---|---|
| Time | $10^{-4}$ $mm^2$/Pa.s | $10^{-5}$ $mm^2$/Pa.s | $10^{-8}$ $mm^2$/Pa.s | $10^{-10}$ $mm^2$/Pa.s | $10^{-11}$ $mm^2$/Pa.s | $10^{-14}$ $mm^2$/Pa.s |
| 5 | 0.003 | 0.003 | 0.002848 | 0.000402 | 4.33E-05 | 4.36E-08 |
| 5.1 | 1.59E-07 | 1.59E-06 | 0.000973 | 0.000375 | 4.09E-05 | 4.13E-08 |
| 5.38 | 1.73E-09 | 1.76E-08 | 0.000157 | 0.000312 | 3.52E-05 | 3.57E-08 |
| 5.804 | 1.61E-09 | 1.61E-08 | 2.97E-05 | 0.000241 | 2.87E-05 | 2.92E-08 |
| 6.3432 | 1.46E-09 | 1.46E-08 | 1.56E-05 | 0.000183 | 2.30E-05 | 2.35E-08 |
| 6.97456 | 1.30E-09 | 1.30E-08 | 1.30E-05 | 0.000141 | 1.87E-05 | 1.93E-08 |
| 7.67965 | 1.13E-09 | 1.13E-08 | 1.14E-05 | 0.000114 | 1.58E-05 | 1.64E-08 |
| 8.44372 | 9.75E-10 | 9.75E-09 | 9.85E-06 | 9.74E-05 | 1.40E-05 | 1.46E-08 |
| 9.25497 | 8.32E-10 | 8.32E-09 | 8.45E-06 | 8.73E-05 | 1.30E-05 | 1.36E-08 |
| 10.104 | 7.04E-10 | 7.04E-09 | 7.18E-06 | 8.08E-05 | 1.24E-05 | 1.30E-08 |
| 10.9832 | 5.91E-10 | 5.91E-09 | 6.07E-06 | 7.63E-05 | 1.21E-05 | 1.27E-08 |
| 11.8865 | 4.94E-10 | 4.94E-09 | 5.10E-06 | 7.28E-05 | 1.19E-05 | 1.26E-08 |
| 12.8092 | 4.11E-10 | 4.11E-09 | 4.27E-06 | 6.98E-05 | 1.18E-05 | 1.25E-08 |
| 13.7474 | 3.41E-10 | 3.41E-09 | 3.57E-06 | 6.71E-05 | 1.17E-05 | 1.25E-08 |
| 14.6979 | 2.82E-10 | 2.82E-09 | 2.97E-06 | 6.46E-05 | 1.16E-05 | 1.24E-08 |
| 15.6583 | 2.33E-10 | 2.33E-09 | 2.46E-06 | 6.21E-05 | 1.15E-05 | 1.24E-08 |
| 16.6267 | 1.92E-10 | 1.92E-09 | 2.04E-06 | 5.98E-05 | 1.15E-05 | 1.24E-08 |
| 17.6013 | 1.58E-10 | 1.58E-09 | 1.69E-06 | 5.75E-05 | 1.14E-05 | 1.24E-08 |

| | | | | | | |
|---|---|---|---|---|---|---|
| 18.5811 | 1.30E-10 | 1.30E-09 | 1.40E-06 | 5.53E-05 | 1.14E-05 | 1.24E-08 |
| 19.5649 | 1.07E-10 | 1.07E-09 | 1.16E-06 | 5.32E-05 | 1.13E-05 | 1.24E-08 |
| 20.5519 | 8.74E-11 | 8.74E-10 | 9.55E-07 | 5.11E-05 | 1.13E-05 | 1.24E-08 |
| 21.5415 | 7.17E-11 | 7.17E-10 | 7.89E-07 | 4.92E-05 | 1.12E-05 | 1.24E-08 |
| 22.5332 | 5.88E-11 | 5.88E-10 | 6.51E-07 | 4.73E-05 | 1.12E-05 | 1.24E-08 |
| 23.5266 | 4.82E-11 | 4.82E-10 | 5.37E-07 | 4.54E-05 | 1.11E-05 | 1.24E-08 |
| 24.5212 | 3.95E-11 | 3.95E-10 | 4.43E-07 | 4.37E-05 | 1.11E-05 | 1.24E-08 |
| 25.517 | 3.23E-11 | 3.23E-10 | 3.65E-07 | 4.20E-05 | 1.10E-05 | 1.24E-08 |
| 26.5136 | 2.65E-11 | 2.65E-10 | 3.01E-07 | 4.03E-05 | 1.10E-05 | 1.24E-08 |
| 27.5109 | 2.17E-11 | 2.17E-10 | 2.48E-07 | 3.88E-05 | 1.09E-05 | 1.24E-08 |
| 28.5087 | 1.77E-11 | 1.77E-10 | 2.04E-07 | 3.73E-05 | 1.09E-05 | 1.24E-08 |
| 29.507 | 1.45E-11 | 1.45E-10 | 1.68E-07 | 3.58E-05 | 1.08E-05 | 1.24E-08 |
| 30.5056 | 1.19E-11 | 1.19E-10 | 1.39E-07 | 3.44E-05 | 1.08E-05 | 1.24E-08 |
| 31.5045 | 9.73E-12 | 9.73E-11 | 1.14E-07 | 3.31E-05 | 1.07E-05 | 1.24E-08 |
| 32.5036 | 7.97E-12 | 7.97E-11 | 9.41E-08 | 3.18E-05 | 1.07E-05 | 1.24E-08 |
| 33.5029 | 6.52E-12 | 6.52E-11 | 7.75E-08 | 3.05E-05 | 1.06E-05 | 1.24E-08 |
| 34.5023 | 5.34E-12 | 5.34E-11 | 6.39E-08 | 2.93E-05 | 1.06E-05 | 1.24E-08 |
| 35.5018 | 4.37E-12 | 4.37E-11 | 5.26E-08 | 2.82E-05 | 1.05E-05 | 1.24E-08 |
| 36.5015 | 3.57E-12 | 3.57E-11 | 4.33E-08 | 2.71E-05 | 1.05E-05 | 1.24E-08 |
| 37.5012 | 2.92E-12 | 2.92E-11 | 3.57E-08 | 2.60E-05 | 1.04E-05 | 1.24E-08 |
| 38.5009 | 2.39E-12 | 2.39E-11 | 2.94E-08 | 2.50E-05 | 1.04E-05 | 1.24E-08 |
| 39.5007 | 1.96E-12 | 1.96E-11 | 2.42E-08 | 2.40E-05 | 1.03E-05 | 1.24E-08 |
| 40.5006 | 1.60E-12 | 1.60E-11 | 1.99E-08 | 2.31E-05 | 1.03E-05 | 1.24E-08 |
| 41.5005 | 1.31E-12 | 1.31E-11 | 1.64E-08 | 2.22E-05 | 1.02E-05 | 1.24E-08 |
| 42.5004 | 1.07E-12 | 1.07E-11 | 1.35E-08 | 2.13E-05 | 1.02E-05 | 1.24E-08 |
| 43.5003 | 8.78E-13 | 8.78E-12 | 1.11E-08 | 2.05E-05 | 1.01E-05 | 1.24E-08 |
| 44.5002 | 7.19E-13 | 7.19E-12 | 9.18E-09 | 1.97E-05 | 1.01E-05 | 1.24E-08 |
| 45.5002 | 5.88E-13 | 5.88E-12 | 7.56E-09 | 1.89E-05 | 1.00E-05 | 1.24E-08 |
| 46.5002 | 4.81E-13 | 4.81E-12 | 6.23E-09 | 1.81E-05 | 1.00E-05 | 1.24E-08 |
| 47.5001 | 3.94E-13 | 3.94E-12 | 5.13E-09 | 1.74E-05 | 9.95E-06 | 1.24E-08 |
| 48.5001 | 3.22E-13 | 3.22E-12 | 4.23E-09 | 1.67E-05 | 9.90E-06 | 1.24E-08 |
| 49.5001 | 2.64E-13 | 2.64E-12 | 3.48E-09 | 1.61E-05 | 9.86E-06 | 1.24E-08 |
| 50.5001 | 2.16E-13 | 2.16E-12 | 2.87E-09 | 1.55E-05 | 9.81E-06 | 1.24E-08 |
